# Wild yeast isolation by middle school students reveals features of North American oak populations of *Saccharomyces cerevisiae* and *Kluyveromyces lactis*

**DOI:** 10.1101/2024.06.27.601111

**Authors:** Randi Yeager, Lydia Heasley, Nolan Baker, Vatsal Shrivastava, Julie Woodman, Michael McMurray

## Abstract

Features of the natural life cycle of the budding yeast *Saccharomyces cerevisiae* were crucial to its domestication as a laboratory experimental model, especially the ability to maintain stable haploid clones and cross them at will to combine alleles via meiosis. Stable haploidy results from mutations in *HO*, which encodes an endonuclease required for haploid-specific mating-type switching. Previous studies found an unexpected diversity of *HO* alleles among natural isolates within a small geographic area. We developed a hands-on field and laboratory activity for middle school students in Denver, Colorado, USA to isolate wild yeast from oak bark, identify species via DNA sequencing, and sequence *HO* from *S. cerevisiae* isolates. We find limited *HO* diversity in North American oak isolates, pointing to efficient, continuous dispersal across the continent. By contrast, we isolated the “dairy yeast”, *Kluyveromyces lactis*, from a tree <10 m away and found that it represents a new population distinct from an oak population in an adjacent state, pointing to high genetic diversity. The outreach activity partnered middle school, high school, and university students in making scientific discoveries and can be adapted to other locations and natural yeast habitats. Indeed, a pilot sampling activity in southeast Texas yielded *S. cerevisiae* oak isolates with a new allele of *HO* and, from a nearby prickly pear cactus, a heat-tolerant isolate of *Saccharomyces paradoxus*.

## INTRODUCTION

Despite an overall excess of workers seeking science, technology, engineering, and mathematics (STEM) careers, women and ethnic and racial minorities remain underrepresented in the academic sector of STEM fields in the United States (Fry *et al*., 2021). Middle school (grades 6-8 in the United States, typically ages 11-14) represents an important educational period in which interest in science as a career is either cultivated or lost (Tai *et al*., 2006). Recent research has demonstrated that informal, extracurricular experiential STEM-related activities can increase interest in future STEM careers among underrepresented populations (Maiorca *et al*., 2021). We sought to develop and test such an activity focused on biology and derived from our research experience with the budding yeast *Saccharomyces cerevisiae*.

Fermentation by *S. cerevisiae* to produce ethanol, carbon dioxide, and flavorful compounds has been exploited by humans for millennia. In the last century *S. cerevisiae* has also moved to the forefront of experimental molecular biology (Botstein and Fink, 2011), due in large part to efficient genetic manipulation coupled with rapid cell proliferation. The last decade has seen a surge in studies of the natural history of *S. cerevisiae* in the hopes of developing a better appreciation of how this experimental workhorse normally lives in the wild (Liti, 2015). Early investigations into the *S. cerevisiae* life cycle funded in part by beer industries in Denmark and the United States (Winge, 1935; Lindegren, 1945) revealed that while diploid yeast typically proliferate via budding (i.e., vegetatively), when starved for nitrogen most natural isolates are able to undergo meiosis accompanied by sporulation to produce haploid spores encased in specialized, stress-resistant cell walls. Without nutrients, a spore remains dormant and for weeks or longer retains the ability to germinate upon restoration of nutrients and either mate with an adjacent haploid cell or proliferate by budding (Brengues *et al*., 2002; Maire *et al*., 2020). Mating occurs only between haploid cells of compatible mating types, **a** with ⍺. Most natural *S. cerevisiae* isolates are diploid (Peter *et al*., 2018), and in most cases a haploid spore of either haploid mating type is able to switch mating types (Fischer *et al*., 2021). Subsequent mating with a daughter cell of opposite mating type provides an efficient route for an isolated spore to return to diploidy.

A major advantage for experimental genetic analysis in *S. cerevisiae* is the ability to stably propagate haploid strains and mate them with other haploids under controlled conditions to produce diploid strains with desired genotypes. A key advance in this regard was the isolation of strains unable to switch mating types (“heterothallic”) (Lindegren and Lindegren, 1943) due to an apparent loss-of-function allele of the *HO* gene (Roberts and Winge, 1949). In strains capable of mating-type switching (“homothallic”), *HO* encodes an endonuclease that makes a single cut in the genome to initiate recombination-mediated exchange of alleles at the *MAT* locus (Strathern *et al*., 1982; Kostriken *et al*., 1983). Apparent loss-of-function *ho* alleles have been found frequently in other natural isolates (Katz Ezov *et al*., 2010; Fischer *et al*., 2021) and in strains used in wine-making (Mortimer, 2000). Since *HO* is not expressed in diploid cells (Jensen *et al*., 1983), any strain that is incapable of, or rarely undergoes, meiosis and/or sporulation would rarely or never express *HO*. Since *HO* is only expressed after a haploid cell buds once (Nasmyth, 1983), *HO* also remains repressed if spores mate immediately upon germination, which is common in some natural isolates (McClure *et al*., 2018). In either scenario the gene could accumulate mutations due to genetic drift in the absence of purifying selection.

Unexpectedly, however, the same alleles are found in isolates from geographically distant locations, inconsistent with drift in the absence of purifying selection. For example, the same *ho* allele originally found near Merced, California was also found in Mount Carmel, Israel, >10,000 km away (Katz Ezov *et al*., 2010). Ho evolved from a domesticated “selfish” genetic element, a homing endonuclease (Coughlan *et al*., 2020). These observations suggest that the common ancestors of isolates with shared *ho* alleles were widely dispersed across the planet, although we cannot exclude the possibility that some residual, ancestral Ho function apart from mating-type switching drove convergent evolution to arrive independently at the same alleles.

For “domesticated” strains associated with human activity, such dispersal could have been the work of humans, as previously suggested for the Merced strain (Katz Ezov *et al*., 2010), but many isolates with shared alleles come from primeval forests or other areas with little human activity (Wang *et al*., 2012) (see Figure 1). While the presence of primeval forest lineages in the bark of planted chestnut trees has been taken as evidence of human involvement in migration (Wang *et al*., 2012), the geographical distribution of *ho* alleles presents an unsolved puzzle. We note that in the age of recreational genetic ancestry testing, the general public is increasingly familiar with the idea of using gene sequences to infer relatedness between geographically distant individuals, yet many people fail to recognize practical applications, such as how genetic testing can prevent disease (Krakow *et al*., 2017). Thus the fundamental concepts underlying this puzzle are relatively straightforward to explain, and present an opportunity to increase genetic literacy with possible implications for future improvements in public health (Abrams *et al*., 2015; Little *et al*., 2022).

**Figure 1.**
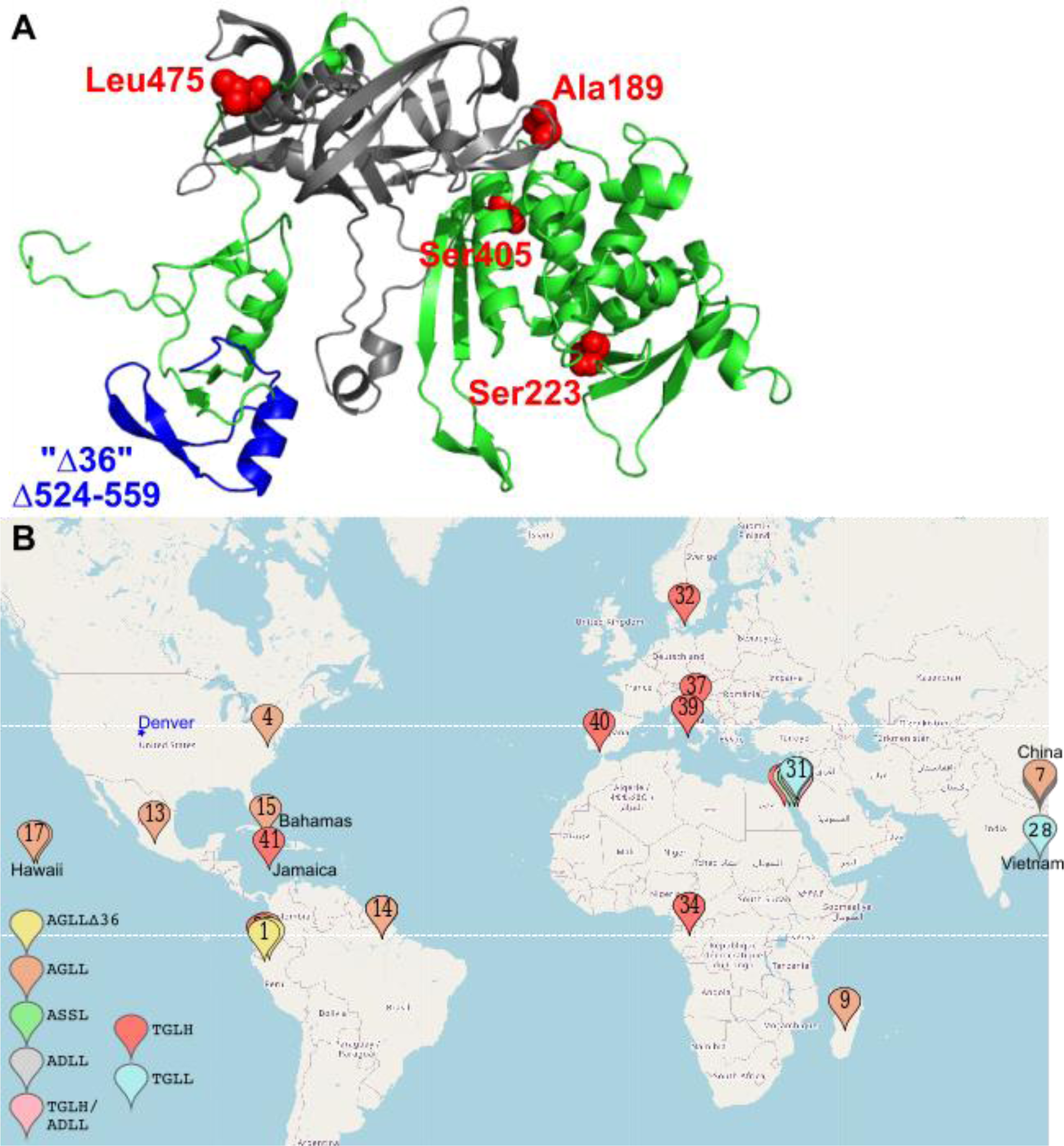
Geographic distribution of shared *ho* alleles among *S. cerevisiae* isolates from nature. (A) Predicted structure of the Ho protein from the S288C genetic background (heterothallic) showing the locations of the regions used for the *HO*-sequence-based isolate grouping in (B): four single amino acid positions and one 36-residue region corresponding to residues 524-559 that is deleted in some isolates (“Δ36”). As delineated by Haber and Wolfe (Haber and Wolfe, 2005) the protein splicing domain is gray. The 36 residues deleted in the “Δ36” deletion are in blue. The structure is oriented so that the parts of the protein that bind DNA are at the bottom. (B) Location of origin for 41 strains of *S. cerevisiae* from the 1,011 genomes collection annotated in that study (Peter *et al*., 2018) as coming from “nature”. Strains were grouped according to *HO* gene sequences corresponding to the regions highlighted in (A). Key indicates genotypes by single-letter amino acid code for the five positions of interest in the order 189, 223, 405, 475 and, if present, the deletion starting at position 524. Numbers on location markers correspond to identifiers in Supplementary Table S1 (see workbook “for MapCustomizer”). Map positions are approximate and some location pins were slightly repositioned to reveal other pins otherwise obscured underneath or to show distant locations on this partial world map. Denver, the site of our isolation activities, is marked with a star.

*Kluyveromyces lactis* is a less familiar yeast species for which there are also interesting questions about the geographical distribution of important genes. Recent studies provided evidence that ancestral strains were unable to ferment lactose, and lactose utilization arose multiple times independently due to strong selection imposed by dairy farmers and the acquisition of a cluster of genes (encoding the galactosidase Lac4 and the lactose permease Lac12) from a closely related species, *Kluyveromyces marxianus* (Friedrich *et al*., 2023). *K. lactis* isolates not associated with human activity lack the gene cluster and utilize lactose less efficiently (Friedrich *et al*., 2023). Genotypic and phenotypic characterization of new *K. lactis* isolates from non-industrial sources may provide additional insights into the geographical distribution of genes associated with lactose utilization. *K. lactis* thus provides a simple illustration of the concept that acquisition of a handful of genes can transform a “wild” organism into an industrially valuable resource, an idea that is broadly accessible to people with basic science knowledge.

We sought to develop a hands-on science outreach activity for middle-school students that addresses the question of how geographically disparate *S. cerevisiae* strains share common *ho* alleles. To broaden the impact of this activity, we further sought to involve university students pursuing Early Education degrees in both carrying out the hands-on activity and developing and evaluating assessments. As a template for others to adapt for their own purposes, here we describe the hands-on activities and present the biological insights derived from our initial efforts with middle-school and university students in the Denver, Colorado Metropolitan region. Pedagogical goals and assessments will be described in detail elsewhere. In addition to basic yeast species identification and *HO* sequence determination for *S. cerevisiae* isolates, we report genotypic and phenotypic characterization of both a natural isolate of *K. lactis* isolated during the outreach activity, and of a heat-tolerant isolate of *Saccharomyces paradoxus*, providing additional research topics readily accessible for outreach activities at the middle-school level.

## MATERIALS AND METHODS

### Oak bark isolation and culture

Sani-cloth Bleach germicidal disposable wipes (PDI # P54072) were used to sterilize the gripping surfaces of six-inch slip-joint pliers (Stanley # 84-097) just prior to bark isolation. The bark fragments were dropped into sterile 50-mL conical tubes. For bark collection in Texas, fragments were dropped into zip lock bags. In the lab, either 10-25 mL of sterile YPD+pen/strep (2% peptone, 1% yeast extract, 2% glucose, 100 units/mL penicillin, 100 µg/mL streptomycin; for the 2018 pilot) or 25 mL of Sniegowski’s enrichment medium (Sniegowski *et al*., 2002) was added to each tube, to submerge the bark. The penicillin and streptomycin were added to the YPD from a 100X stock (HyClone). The resulting cultures were maintained at room temperature (2018 pilot) or 30°C (2019 activity with CCU participation) without constant agitation; they were agitated occasionally, during periodic inspection for turbidity. Three-µL aliquots were removed and placed between a glass slide and #1 thickness 18 mm x 18 mm coverslip for examination by light microscopy with objectives of various magnifications (40X, 60X, or 63X). Images in the figures here were taken with an EVOSfl all-in-one epifluorescence microscope (Thermo-Fisher Scientific) with a 60X oil objective.

For YPD+pen/strep cultures that contained yeast-form cells, ∼500 µL were used to inoculate 25 mL of Sniegowski’s enrichment medium (per L: 3 g yeast extract, 3 g malt extract, 5 g peptone, 10 g sucrose, 76 mL ethanol, 400 µg chloramphenicol, 1 mL 1M hydrochloric acid (Sniegowski *et al*., 2002)) in new 50-mL conical tubes. These cultures were also maintained at room temperature without constant agitation and inspected with a microscope in the same way. For Sniegowski’s cultures with yeast-form cells, aliquots were either diluted in sterile water and spread on solid YPD medium (2% agar) using sterile glass beads, or 3 µL of undiluted culture was spotted on a YPD plate and then the cells were streaked across the rest of the surface of the plate using a sterile toothpick. These plates were incubated at 30°C until colonies were visible. Cells from these colonies were scraped with a pipette tip and resuspended in 3 µL of water on a microscope slide before applying a glass coverslip and imaging.

Colonies composed of cells resembling *S. cerevisiae* were used for sporulation tests by scraping cells from the colony with a sterile toothpick and spreading on a sporulation plate (1% potassium acetate, 2% agar) in an area ∼1 cm^2^. Following incubation at 30°C for 1-2 days, cells from the sporulation plate were scraped off with a pipette tip and imaged. For those that harbored asci resembling *S. cerevisiae* (see Figure 2B), more cells were scraped from the sporulation plate and resuspended in 50 µL of zymolyase (Zymo Research #E1004) diluted 1:50 in water to digest ascus walls. Following ∼10 min of digestion, 3 µL was spread in a strip on the surface of a YPD plate, and a tetrad dissection microscope was used to isolate individual spores. The YPD plate was incubated at 30°C until colonies formed. To test for homothallism, cells from individual colonies were spread on another sporulation plate, and the subsequent appearance of asci was taken as evidence of the presence of diploid cells, indicative of mating-type switching.

**Figure 2.**
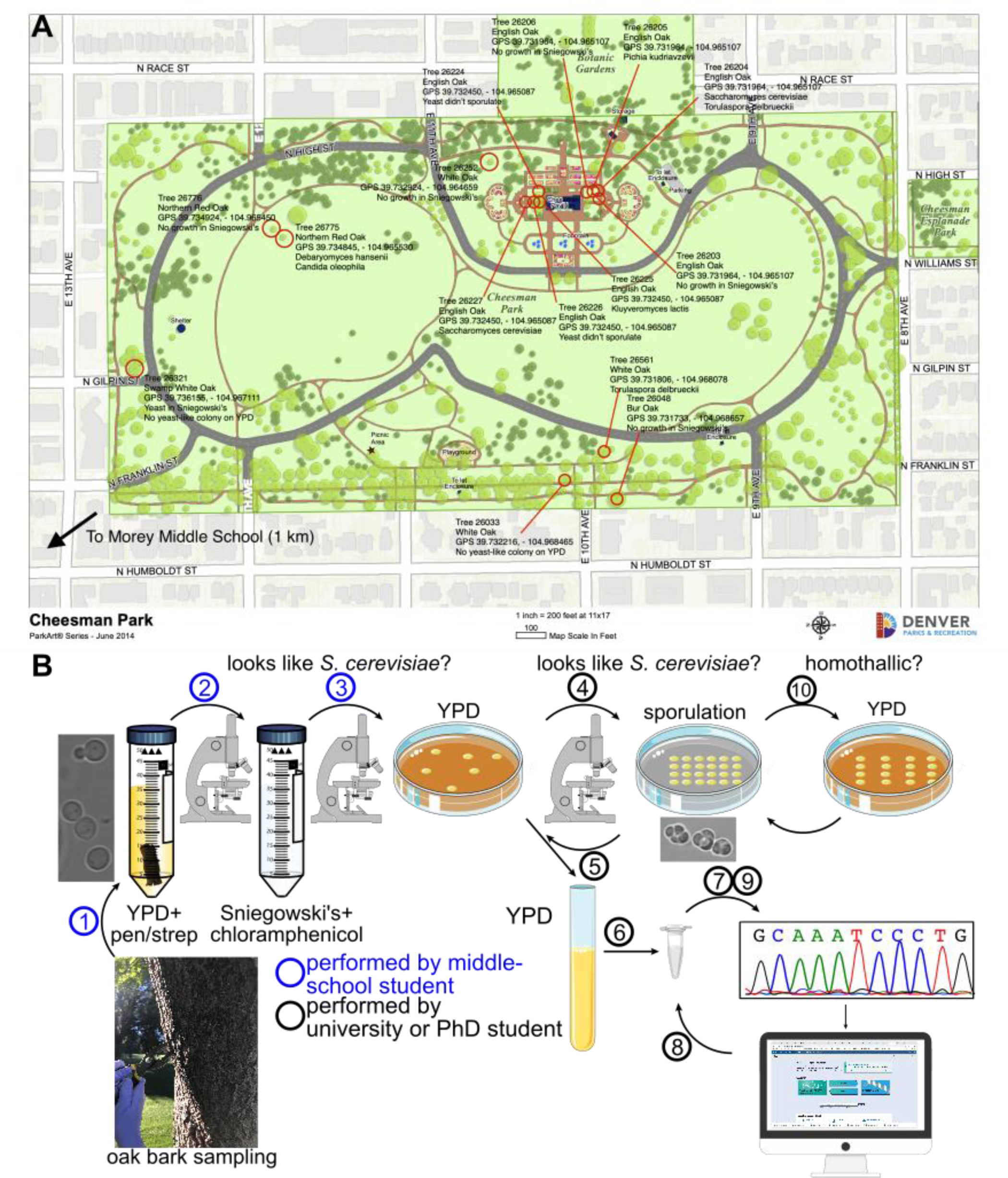
Tree map and schematic illustration of the outreach activity. (A) Map of Cheesman Park with target trees from 2018 pilot activity labeled and annotated with species of yeast isolated, if any. (B) Circled numbers indicate the following steps in the wild yeast isolation activity. (1) Oak bark samples are incubated in liquid rich medium (YPD) with anti-bacterial antibiotics (penicillin and streptomycin, “pen/strep”). (2) After microscopic observation of YPD+pen/strep cultures, aliquots of those cultures that contained yeast-form cells were used to inoculate a more selective enrichment medium containing a different anti-bacterial antibiotic (Sniegowski’s with chloramphenicol). The micrograph at left shows an example image from a YPD+pen/strep culture from tree 26227, from which *S. cerevisiae* was ultimately isolated. (3) Microscopic examination of the selective cultures was used to identify those containing yeast-form cells, aliquots of which were plated to YPD agar plates to allow colonies to form. (4) Cells from individual colonies were examined with a microscope to identify those with yeast-form cells, portions of which were transferred to solid sporulation-inducing medium (1% potassium acetate). (5) Strains that generated asci resembling *S. cerevisiae* (the micrograph shows a representative isolate from tree 26227) were selected for inoculation from the YPD plate into liquid YPD medium. (6) Extraction of genomic DNA was followed by (7) PCR amplification of *ITS2* sequences and subsequent Sanger sequencing. (8) Isolates identified as *S. cerevisiae* by BLAST analysis were further analyzed by PCR and (9) sequencing of the *HO* gene. The chromatograph shows actual *HO* sequence data from an *S. cerevisiae* isolate from tree 26204. (10) *S. cerevisiae* isolates were tested for homothallism by dissecting tetrad asci and allowing spores to germinate and form colonies on YPD. Cells from the resulting colonies were transferred to sporulation-inducing medium and examined for the presence of asci, indicative of mating-type switching after spore germination and mating to return to diploidy.

### Laboratory and reference yeast strains

For PFGE, we used *S. cerevisiae* strain FY2742 (S288C derivative MAT⍺ *his3Δ1 leu2Δ0 lys2Δ0 ura3Δ0 RME1 TAO3 MKT1*, where the alleles at *RME1*, *TAO3* and *MKT1* are from the SK1 strain background (Kloimwieder and Winston, 2011) and prototrophic wild-type diploid *S. paradoxus* strain N17 (Naumov *et al*., 1990). For lactose utilization, we used an *S. cerevisiae* isolate from tree 26204 (Table 1) that we named MMY0374 and *K. lactis* NRRL Y-1140 ATCC #8585 (Dujon *et al*., 2004; Zivanovic *et al*., 2005).

**Table 1.** Oak trees sampled and yeast species isolated in the fall 2018 outreach activity.

| Oak (species, number, GPS coordinates) | YPD+pen/strep yeast | Sniegowski's yeast | Yeast-like colony | Sporulation | ITS species |
| --- | --- | --- | --- | --- | --- |
| *Quercus alba*, 26033, 39.732216, -104.968465 | + | + | – | ND | ND |
| *Quercus robur*, 26203, 39.731964, -104.965107 | ND | + | – | ND | ND |
| *Quercus robur*, 26204, 39.731964, -104.965107 | +, + | +, + | +, + | +, ? | *S. cerevisiae*, *Torulaspora delbrueckii* |
| *Quercus robur*, 26205, 39.731992, -104.965094 | ND, ND | +, + | –, + | ND, – | *Pichia kudriavzevii* |
| *Quercus robur*, 26206, 39.732035, -104.965094 | +, ND | +, + | +, – | +, ND | ND, ND |
| *Quercus robur*, 26224, 39.732450, -104.965087 | ND, + | +, + | +, + | +, + | ND, ND |
| *Quercus robur*, 26225, 39.732450, -104.965087 | +, + | +, + | +, + | +, + | *K. lactis*, ND |
| *Quercus robur*, 26226, 39.732450, -104.965087 | ND, ND | +, + | –, + | ND, + | ND, ND |
| *Quercus robur*, 26227, 39.732450, -104.965087 | +, ND | +, + | + (x3), + | +, + | *S. cerevisiae* (x2), unknown |
| *Quercus alba*, 26252, 39.732924, -104.964659 | ND | + | + | ? | *Torulaspora delbrueckii* |
| *Quercus bicolor*, 26321, 39.736155, -104.967111 | ND | + | – | ND | ND |
| *Quercus alba*, 26561, 39.731806, -104.968078 | ND | + | + | ? | *Torulaspora delbrueckii* |
| *Quercus rubra*, 26775, 39.734859, -104.965500 | ND | + | + (x2) | ?, ND | *Debaryomyces hansenii*, *Candida oleophila* |

ND, not determined.
+, cells or colony with yeast-like appearance were detected
–, no cell or colony with yeast-like appearance was detected
?, ambiguous/unclear results
(x2), two distinct colonies or isolates were found.

### DNA isolation, PCR, and single-gene sequencing

Genomic DNA was isolated from cells pelleted from 5-mL overnight YPD liquid cultures or scraped from YPD plates using a modified *E. coli* plasmid DNA isolation protocol/kit (Johnson *et al*., 2015). Standard Taq (New England Biolabs #M0273) or Phusion (New England Biolabs #M0530) or Q5 (New England Biolabs #M0491) DNA polymerase was used for PCR according to the manufacturer’s instructions, with 1 µL of genomic DNA as template. PCR for species identification (White, 1990) used primers ITS1 (TCCGTAGGTGAACCTGCGG), ITS4 (TCCTCCGCTTATTGATATGC), NL1 (GCATATCAATAAGCGGAGGAAAAG), and/or NL4 (TCCTCCGCTTATTGATATGC). PCR for *HO* sequencing used primers 5’ScereHOfw (TCCATATCCTCATAAGCAGC), ScHOmidre (CCTGCGAGTACTGGACCAAA), S.cereHOmidfw (CAGCTACTGTGACGACCAGG), HOmidfw2 (GGTCTGTGGTTAGGTGACAG), HOmidre2 (TCTCACCCTGGAAATCATCC), HOendrev (TACCACAACTCTTATGAGGC), and/or 3’HOre (TGGCGTATTTCTACTCCAGC). After inactivation/digestion of primers and nucleotides via treatment with Exonuclease I (Thermo Scientific Fermentas #EN0581) and alkaline phosphatase (Thermo Scientific Fermentas #EN0581), the products were sequenced via Sanger sequencing with the same primers. Contigs were assembled using the CAP3 Sequence Assembly Program (https://doua.prabi.fr/software/cap3) (Huang and Madan, 1999).

For one amplicon from genomic DNA of MMY0358 using the primers 5’ScereHOfw and HOmidre2, the PCR product was purified using the DNA Clean & Concentrator-5 kit (Zymo Research #D4013) and direct long-read sequencing was performed by Quintara Biosciences, Inc.

### Maximum temperature estimation

We viewed daily temperature data for Denver from the months of June, July, and August of 2012-2022 at https://www.weather.gov/bou/local_climate and averaged the values for the highest daily temperature recorded for each month. No data were available for July 2015. For College Station, Texas, we used online weather data at https://www.weather.gov/wrh/Climate?wfo=hgx and browsed the monthly highest max temperature from 2012-2019 at Easterwood Field, College Station.

### Phylogenetic analysis

The *K. lactis ACT1* phylogenetic tree was based on nucleotides 819-1904 where position 1 is the start codon. *S. cerevisiae ACT1* sequences were aligned using the Simple Phylogeny tool at the EMBL-EBI server (www.ebi.ac.uk/Tools/phylogeny/simple_phylogeny/) from a Clustal Omega alignment performed at the same server. Input Parameters were: Tree Format: phylip, No distance correction, gaps were not excluded, and a Neighbour-joining Clustering Method was used.

### Pulsed-field gel electrophoresis

Yeast strains were grown in 7 mL liquid YPD cultures at 30°C for 24 hours. Cells were pelleted and resuspended in low melting point agarose to form plugs. Plugs were treated with zymolyase to remove cell walls and proteinase K to degrade protein components. Plugs were loaded into a 1% gel made in tris boric acid EDTA buffer (45 mM Tris-borate 1 mM EDTA). The gel was then run in the same buffer in a BioRad CHEF DRII Mapper for 60 hours with initial switch and final switch times 50s and 150s, respectively. To visualize size-resolved chromosomes, the gel was stained with ethidium bromide.

### Lactose utilization assay

5-mL YPD liquid cultures were inoculated with colonies of Y0329, Y0374, or NRRLY-1140 from YPD plates and rotated at 30°C overnight. 1 mL of each culture was pelleted in a microcentrifuge tube by a pulse of centrifugation at maximum speed. The medium was removed and the cells were resuspended in 1 mL sterile distilled deionized water, then pelleted again. After resuspending the cells again in 1 mL water, 100 µL of each was mixed with 15 mL of lactose (per L: 6.7 g yeast nitrogen base with amino acids and ammonium sulfate (Research Products International # Y20030, 100 g lactose) or glucose (per L: 6.7 g yeast nitrogen base with amino acids and ammonium sulfate, 20 g glucose) medium in a petri dish, then 150 µL of this suspension was transferred with a multichannel pipette to each of 16 wells of a flat-bottom 96-well plate. The plate was incubated for 22 hr 9 min at 30°C in a BioTek Cytation 3 plate reader and the absorbance at 595 nm was read every 5 minutes following orbital shaking. Eight wells away from the plate edges were used for analysis.

### Comparing culture growth rates at different temperatures

Yeast cells grown on a solid YPD plate were resuspended in liquid YPD in sterile glass culture tubes, from which 100 µL was transferred to 8 wells of a 96-well plate of PCR tubes that was cut in half. The plate was incubated in a thermal cycler using a program with gradient of temperatures over 8 wells from 30°C to 45°C. The lid temp was set to 50°C. After 21 hr incubation, 50 µL of each culture was transferred to the wells of a 96-well plate, after first reading the absorbance at 595 nm of the empty wells using a Cytation 3 plate reader (Bio-Tek). The absorbance of each culture was measured and the values of the empty wells were subtracted.

### Yeast whole genome sequencing

DNA extraction and sequencing was performed by The Sequencing Center, Fort Collins, Colorado. Zymo Research Quick-DNA Fungal/Bacterial Microprep Kit (#D6007) was used to extract DNA from *K. lactis* strain MMY0329. Cells (10-20 mg wet weight) were resuspended in 200 μL phosphate buffered saline (PBS) and added to a ZR BashingBead™ Lysis Tube (0.1 & 0.5 mm) with 750 μL Lysis Solution. The tube was processed in a Qiagen TissueLyser LT bead beater at 50 Hz for 5 min and centrifuged at 10,000 × *g* for 1 min. 400 μL supernatant was transferred to a Zymo-Spin™ IV Spin Filter in a Collection Tube and centrifuged at 7,000 × *g* for 1 min. ß-mercaptoethanol was added to Genomic Lysis Buffer to a final concentration of 0.5% (v/v). 1.2 mL Genomic Lysis Buffer was added to the filtrate. 800 μL of the mixture was transferred to a Zymo-Spin™ IC Column in a Collection Tube and centrifuged at 10,000 × *g* for 1 min. The flow-through was discarded and the previous step was repeated once. 200 uL DNA Pre-Wash Buffer was added to the Zymo-Spin™ IC Column and centrifuged at 10,000 × *g* for 1 min. 500 µL g-DNA Wash Buffer was added to the Zymo-Spin™ IC Column and centrifuged at 10,000 × *g* for 1 min. The Zymo-Spin™ IC Column was transferred to a 2-mL microcentrifuge tube and 20 μL of DNA Elution Buffer was added directly to the column matrix. After 3 min. incubation, the column was centrifuged at 10,000 × g for 30 seconds to elute ultra-pure DNA.

The Illumina Nextera XT DNA Library Prep Kit was used to prepare extracted yeast DNA for sequencing. For tagmentation, 10 μL of Tagment DNA Buffer and 5 μL of extracted gDNA were added to each plate well in a PCR plate and mixed by pipetting. 5 μL of Amplicon Tagment Mix was added to each plate well, mixed by pipetting and the plate was centrifuged at 280 × *g* for 1 min. A predefined tagmentation program was run on an ABI GeneAmp PCR System 9700 thermal cycler. Immediately following its completion, 5 μL of Neutralize Tagment Buffer was added to each plate well and mixed by pipetting. The plate was centrifuged at 280 × *g* for 1 min followed by incubation at room temperature for 5 min. For amplification and indexing, Illumina Index 1 (i7) adapters and Index 2 (i5) adapters were uniquely assigned to each sample on an index adapter plate. 15 μL of NexteraPCR Master Mix (NPM) was added to each plate well and mixed by pipetting. The Index/NPM mixture was transferred to the PCR plate, mixed by pipetting, and centrifuged at 280 × g for 1 min prior to PCR. Library cleanup was performed using AMPure XP beads (Beckman). The PCR product was centrifuged at 280 × *g* for 1 min and 50 μL of product was transferred to a new PCR plate. 30 μL of AMPure XP beads was added to each plate well and the plate was shaken at 1800 rpm for 2 min. The plate was incubated at room temperature for 5 min and placed on a magnetic stand until the liquid was clear. All supernatant was removed and discarded and the beads were washed twice with 200 μL 80% ethanol. The plate was air-dried on a magnetic stand for 15 min. 52.5 μL of RSB was added to the beads and the plate was shaken at 1800 rpm for 2 min. The plate was incubated at room temperature for 2 min and placed on a magnetic stand until the liquid was clear. 50 μL of supernatant was transferred to a new PCR plate. To check library size distribution, undiluted library was analyzed on an Agilent Technologies 2200 TapeStation using Agilent Technologies High Sensitivity D1000 ScreenTape.

44 μL of Library Normalization Additives 1 (LNA1) per sample and 8 uL of Library Normalization Beads 1 (LNB1) per sample were added to a 15-mL conical tube. 45 μL of the LNA1/LNB1 mixture was added to each sample PCR plate well. The plate was shaken at 1800 rpm for 30 min and placed on a magnetic stand until the liquid was clear. The supernatant was removed and discarded. The beads were washed twice with 45 μL of Library Normalization Wash 1 and 30 μL of 0.1 N sodium hydroxide was added to each plate well. The plate was shaken at 1800 rpm for 5 min. After a 5-min incubation, the samples were resuspended and shaken at 1800 rpm for 5 min. The plate was placed on a magnetic stand until the liquid was clear. The supernatant was transferred to a new PCR plate and the plate was centrifuged at 1000 × g for 1 min. 5 μL of each library was transferred from the PCR plate to a 2-mL microcentrifuge tube. A fresh dilution of 0.1 N sodium hydroxide was prepared by combining 900 μL of laboratory-grade water with 100 μL of stock 1.0 N sodium hydroxide. The denaturation process involved combining 5 μL of 1 nM library with 5 μL of 0.1 N sodium hydroxide, vortexing and incubating at room temperature for 5 min. 5 μL of 200 mM Tris-HCl, pH 7.0, was added to the denatured library, vortexed and centrifuged at 280 × *g* for 1 min. The denatured library was diluted with 985 μL Hybridization Buffer, vortexed and centrifuged at 280 × *g* for 1 min. The resulting diluted library was transferred to a 2-mL microcentrifuge tube and further diluted with 320 μL Hybridization Buffer to achieve a final loading concentration of 1.8 pM. A 5% PhiX spike-in control was added to the final library. Yeast whole genome sequencing was performed on an Illumina MiniSeq short-read sequencer using a standard Illumina workflow and configured for 2 x 150 bp paired-end reads and MiniSeq flow cell.

## RESULTS

### Disparate origins of *S. cerevisiae* isolates with shared alleles at *HO*

Others previously noted the diversity of alleles of *HO* among the genome sequences of 1,011 *S. cerevisiae* isolates and the fact that many are predicted to be non-functional for mating-type switching (Peter *et al*., 2018; Fischer *et al*., 2021). We examined the geographical origin of those strains with respect to the *HO* allele of each. We focused on five locations in the *HO* coding sequence, four single residues (189, 223, 405 and 475) where substitutions were found in a heterothallic fig isolate compared to the “wild-type” sequence from homothallic strains (Meiron *et al*., 1995), and one 36-residue region (residues 524-559) previously documented as being absent in a heterothallic isolate from Brazil (Argueso *et al*., 2009). The residue at position 189 is not conserved in related yeast species and lies in a linker between the protein splicing and endonuclease domains of Ho (Figure 1A). Ala, Thr or occasionally Ser is encoded here, with no apparent effect on Ho function. Residue 223 lies in one of two “endonuclease motifs” conserved between Ho and PI-*Sce*I, a closely-related yeast endonuclease (Bakhrat *et al*., 2004), and is Gly in PI-*Sce*I and all known homothallic strains as well as in strains from primeval Chinese forests (e.g. strain BAM) where *S. cerevisiae* is thought to have originally evolved (Wang *et al*., 2012; Peter *et al*., 2018). Ser in this position is sufficient to render an otherwise active allele inactive (Ekino *et al*., 1999). Residue 405 lies in the endonuclease domain but outside conserved regions (Haber and Wolfe, 2005). Leu is found here in many homothallic strains but alleles encoding both Ser in this position and Leu475 show some activity (Ekino *et al*., 1999). Residue 475 lies in one of two zinc finger domains predicted to directly contact DNA, and the H475L substitution was originally proposed to disrupt Ho function (Meiron *et al*., 1995) but a subsequent study found some mating-type switching activity of an allele encoding Leu475 (and Ala189 Gly223 and Ser405) (Ekino *et al*., 1999).

The deleted region is interesting in two ways. First, the deletion rather precisely removes the other zinc finger, hence the deletion allele might bind a different DNA sequence, or none at all. Second, it is flanked by repeats of an 8-nucleotide sequence that likely act to promote homologous recombination that excises the intervening sequences (Argueso *et al*., 2009). Thus the deletion would be expected to occur spontaneously at a higher rate than a point mutation.

To look for shared alleles, we downloaded the *HO* coding sequences from the 1,011 genomes study (http://1002genomes.u-strasbg.fr/) and used Microsoft Excel to search within each for short (7-16-nucleotide) sequences surrounding each of the positions introduced above, then grouped isolates that shared the same sequences for all five positions. Since in the 1,011 genomes mixed sequencing results at a given position are indicated with the standard single-letter codes for multiple nucleotides (e.g., R for A or G), we detected heterozygosity in many cases and some groups shared heterozygosity at specific positions. We ignored the 124 isolates for which *HO* was deliberately deleted in the lab, and focused on the 41 isolates annotated as being of “natural” origin.

As shown in Figure 1B and Supplementary Table S1 for seven *HO* groups, isolates from the same group originated in geographically disparate locations. A phylogenetic tree made from the *HO* sequences recapitulated most of the major relationships established in the 1,011 genomes study using thousands of SNPs across the genome (Figure S1). Satisfyingly, the group heterozygous at residues 189 (Ala/Thr), 223 (Asp/Gly) and 475 (Leu/His) clusters with the 1,011 genomes “mixed origin” clade near the group with Ala189, Asp223, and Leu475 and the group with Thr189, Gly223, and His475 (Figure S1). Mixing between these groups could explain the origins of the two alleles in the heterozygous strains. Both strains with the 36-residue deletion (ALC and ALH, isolated from distinct locations in Ecuador) also encode a single-nucleotide deletion at codon 138 resulting in a predicted truncation after 148 residues (Figure S1B). The same frameshift is also found in three strains from Mount Carmel (BDC, BDF, and BDG) that lack the 36-residue deletion but otherwise differ by only three nucleotides from the Ecuadorian strains (Supplementary Table S1). If the frameshift eliminates Ho function, it is easy to imagine how the 36-residue deletion could subsequently arise and persist by avoiding purifying selection. How isolates from Ecuador and Israel so closely resemble each other is more mysterious.

A subsequent study of the 1,011 *S. cerevisiae* isolates assessed sporulation ability (De Chiara *et al*., 2022). There was no obvious correlation among natural isolates between any specific allele of *HO* and the ability to sporulate efficiently (Supplementary Table S1, see workbook “Natural isolates”). These observations do not support our speculation (see Introduction) that a failure to sporulate in the wild is the source of allelic variation in *HO* because it precludes the ability of purifying selection to maintain specific (functional) *HO* alleles.

### Engaging middle-school students in isolation of wild yeast from oak bark

We sought out middle school students near our institutions to develop an activity engaging those students in the search for natural *S. cerevisiae* isolates and *HO* alleles. Morey Middle School in Denver, Colorado is an urban school serving sixth- to eighth-graders in the Denver Public School district. We contacted the sixth grade Science teacher at Morey at the beginning of the 2018/2019 and 2019/2020 school years to determine when and how our activity might best fit with her planned curriculum, and further designed our activity to meet Colorado middle school academic standards and learning expectations for Life Science (Supplementary Table S2). A ∼40-minute didactic lecture presentation in the classroom, accompanied by image- and video-rich slides, introduced the specific topics. Altogether, the didactic material and the hands-on activity (see below) match six of twelve Colorado state Life Science learning standards (Supplementary Table S2). We chose dates for the lectures and hands-on activity in between the Science classes’ units on Microbiome, Metabolism and Traits and Heredity.

To develop a hands-on activity, we adapted existing protocols developed by others for yeast isolation from the bark of oak trees (Sniegowski *et al*., 2002). Though *Saccharomyces* yeasts are likely minor components (Kowallik *et al*., 2015), oak bark is a known habitat for many yeast species, including *S. cerevisiae* (Naumov *et al*., 1998). Cheesman Park is within walking distance (1 km) of Morey Middle School and is maintained by the City of Denver, which also maintains an interactive online inventory of tree species (TreeKeeper, denverco.treekeepersoftware.com). TreeKeeper and Google Maps (http://www.google.com/maps/) were used to prepare a map of Cheesman Park in which all target oak trees were identified by numbers (from TreeKeeper) and assigned global positioning satellite (GPS) locations using Google Maps. Trees were grouped by threes and assigned to groups of students.

In the 2018 pilot activity, an adult led groups of four to six students to oak trees of a variety of species, where they collected ∼2-cm pieces of bark using kits of nitrile gloves, 50-mL conical tubes pre-filled with 10 mL liquid yeast peptone dextrose medium supplemented with the antibacterial drugs penicillin and streptomycin (“YPD+pen/strep”), pliers, sterilizing wipes, and a laminated tree map (Figure 2A). Bark (one piece per culture) was chosen from parts of the trees on or near their trunks, 0.5 to 1 m from the soil, as a compromise between ease of accessibility and distance from suspected sources of “contamination” (humans and domesticated animals). Bark cultures (two per tree, 15 trees) were incubated in the classroom for several days and then transferred to the McMurray lab. Samples from each culture were examined by transmitted light microscopy. Yeast-form (ovoid) cells were present in most YPD+pen/strep cultures, but non-yeast cells were predominant (Figure 2B). At this point, the students were asked to complete during class an online multiple-choice quiz (prepared in Google Forms; Figure 2B) distinguishing yeast-form cells from others in micrographs from selected cultures.

To enrich for yeast, aliquots of cultures with any visible growth (turbidity) were transferred to Sniegowski’s medium, which has a high ethanol content (Sniegowski *et al*., 2002). 21 Sniegowski’s cultures contained yeast-form cells (Table 1). Dilutions of each were plated to solid YPD medium, from which single colonies were examined for yeast-form cells. 15 resulting yeast strains were examined microscopically after transfer to sporulation medium; six formed asci resembling *Saccharomyces* (Table 1; see Figure 2). Species identification was performed using rDNA internal transcribed spacer (*ITS2*) sequences amplified by PCR from genomic DNA. Sequencing identified three *S. cerevisiae* isolates; seven other strains were a variety of non-*Saccharomyces* yeasts, including *Kluyveromyces lactis* (Table 1). Sequence quality from one strain was too poor for unambiguous species identification, and four were not sequenced (Table 1).

Notably, we did not isolate *Saccharomyces paradoxus*, a species closely related to *S. cerevisiae* and often found together in the same oak forests or even on the same tree (Sniegowski *et al*., 2002; Sampaio and Gonçalves, 2008; Liti *et al*., 2009; Zhang *et al*., 2010). The failure to isolate *Saccharomyces paradoxus* from Denver oaks was not unexpected, since prior research suggested that the geographical distribution of *S. cerevisiae* versus *S. paradoxus* on oak trees can be predicted by the maximum summer temperatures (Robinson *et al*., 2016). Using available local climate data we estimated the average maximum summer temperature for Denver from 2012-2022 as ∼36.75°C, within the maximum temperature range considered optimum for wild *S. cerevisiae* (25-38°C) but too hot for *S. paradoxus* (18-31°C), at least in the oak habitat. These data also agree with recent *S. paradoxus* isolation results from China (He *et al*., 2022).

Homothallism was tested in three of the *S. cerevisiae* isolates by inducing sporulation, isolating individual spores on rich medium, and testing cells within the resulting colonies for the ability to sporulate, as an indicator of the presence of diploid cells. All three isolates sporulated with very high efficiency (see Figure 2). *HO* coding sequences were amplified by PCR and sequenced. All three sequences are of the group with Ala189 Gly223 Leu405 Leu475 and no deletion and are identical to the sequence found in homothallic strains isolated from oak trees in Missouri (“T7” strain) or Pennsylvania, USA (YPS163 strain) (Sniegowski *et al*., 2002; Fay and Benavides, 2005). We note that this *HO* allele must be functional despite encoding Leu405 and Leu475, since YPS163 (Fay and Benavides, 2005) and all our isolates are homothallic.

At this point, about a week after the bark gathering, a final classroom lecture introduced the students to the concept of species identification by sequencing a region of the genome that is sufficiently conserved between species to allow comparison but is sufficiently variable to distinguish between closely related species. The lecture included instructions on how to use the NCBI BLAST server to find the species with the best match. Then, each student was sent a link to access an anonymized *ITS2* sequence from one of the oak bark isolates and asked to perform a BLAST alignment and enter in an online form their best guess as to the species identity. The students were also informed about the homothallism and *HO* sequencing results for the *S. cerevisiae* isolates. This activity concluded the pilot outreach.

The activity was repeated, with some changes, with a new class of middle school students in the 2019-2020 school year. Most important was the inclusion of first-year university students from Colorado Christian University (CCU). Nine Biology majors and 6 Early Education majors were present at Morey for the bark isolation, where they led groups of middle-school students in the activities. Since the Sniegowski’s medium was effective at enriching for yeast, the YPD+pen/strep step was skipped, and bark samples were cultured directly in Sniegowski’s enrichment medium. The cultures were monitored by CCU students in the Woodman lab, and they also tested for sporulation. Additionally, simple light microscopes owned by Morey Middle School were used with microscope slides and coverslips in a hands-on activity in which the students examined the bark cultures in the classroom. A digital eyepiece camera (Levenhuk M300 BASE) connected to a laptop computer was used to project live video from one microscope to a projection screen in the classroom, providing real-time instruction for the students to follow. Finally, the students completed custom assessments to gauge effectiveness of the activity. These assessments and related educational activities will be described elsewhere.

### Geographical distribution of *HO* alleles shared with Denver oak isolates

One new *S. cerevisiae* isolate was obtained, found to be homothallic, and the *HO* sequence was identical to the three from the other Cheesman Park oak isolates. We examined the origins of all 20 isolates from the 1,011 genomes study that share this *HO* sequence. This group includes 12 of the 13 *S. cerevisiae* isolates in the “North American Oak” clade, and four other isolates from North American oaks that were not placed in that clade. One of the isolates is from the “Mosaic region 3” clade, while the remaining four are from human food (from India or Japan) or a North American insect (Supplementary Table S1). Notably, the other isolate from the 1,011 genomes “North American Oak” clade, strain ACD, has an *HO* sequence that differs at six nucleotides and encodes an Ho protein of the “Ala189 Ser223 Ser405 Leu475 and no deletion” group. Its *HO* allele is identical to only four others from the 1,011 genomes, one North American oak isolate (ADH), a human clinical isolate from the USA (ABB), and two wine isolates, one from Russia (ACQ) and one from Ivory Coast (ACK). All other isolates from ostensibly natural North American sources among the 1,011 genomes have different *HO* sequences (Supplementary Table S1). These observations suggest that there is little genetic variation among *S. cerevisiae* residing in North American oaks, and little mixing between North American oak populations and other natural populations on the same continent. Shared sequences with isolates from other continents presumably reflect either human-assisted transportation or an ancient diversity of alleles prior to global distribution. We note that the 2019 isolate came from the same English oak (*Quercus robur*) tree as did one of the *S. cerevisiae* isolates from the 2018 activity. A subsequent isolation activity in 2023 again isolated *S. cerevisiae* from some of the same English oaks, plus other nearby oaks of the same species, and also from two Northern Red oaks (*Quercus rubra*; unpublished results).

### A *K. lactis* isolate from oak bark represents a new natural population

To ask if the lack of genetic diversity of our oak isolates extended to other yeast species, we further characterized the oak bark isolate of *K. lactis* to compare it with the distinct populations of the species that have been established in the literature. Pulsed-field gel electrophoresis (PFGE) of intact chromosomes revealed a small number of large chromosomes (Figure 3A), most similar to previously characterized isolates of a “new” North American genetic population (Lyutova *et al*., 2022). Notably, one member of the “new” North American population was isolated from an oak in Arizona (Naumova *et al*., 2004), a state adjacent to Colorado.

**Figure 3.**
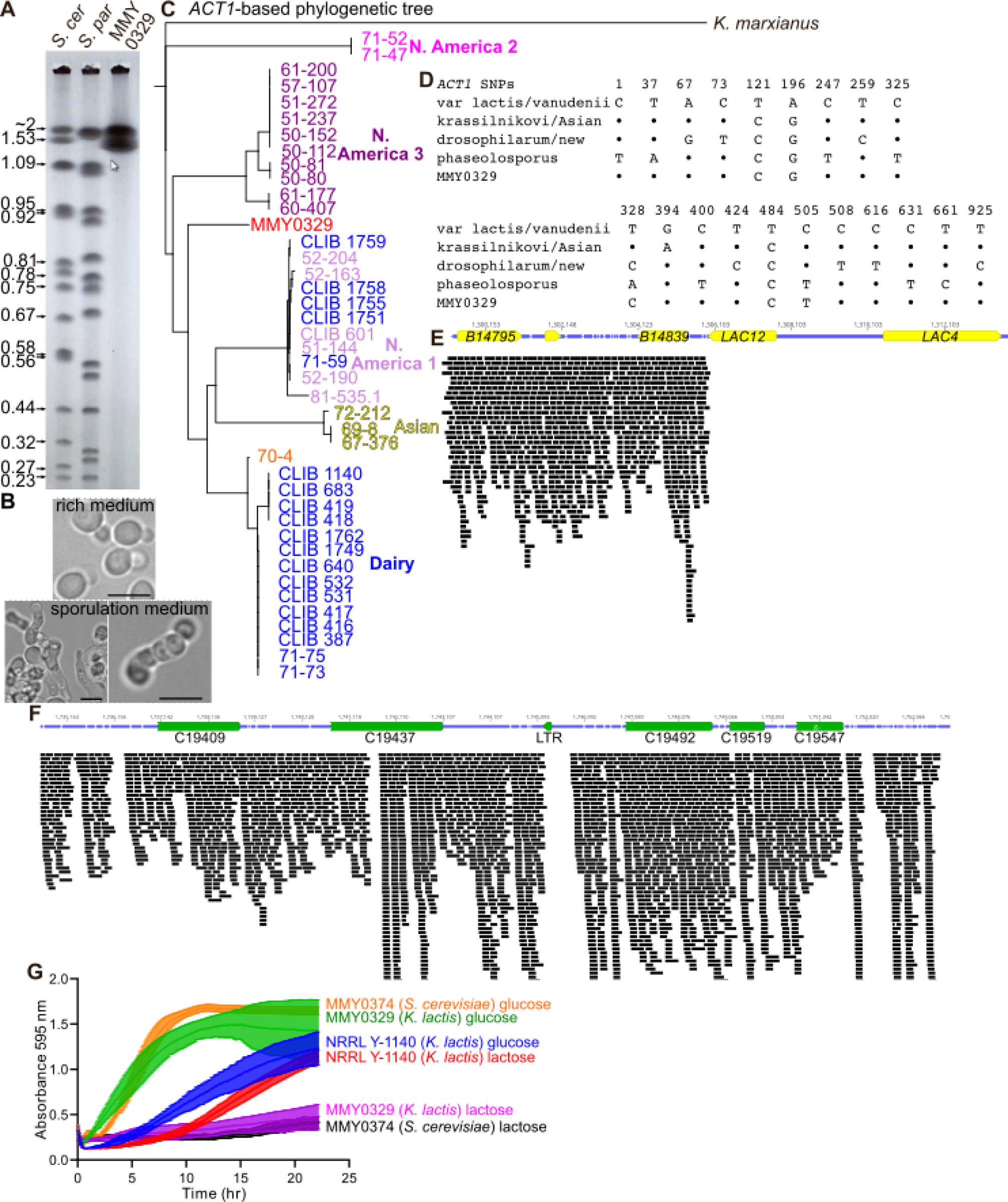
Molecular genetic analysis of a Denver oak isolate of *K. lactis* points to a new population. (A) Pulsed-field gel electrophoresis of intact chromosomes from *S. cerevisiae* (“*S. cer*”, strain FY2742), *S. paradoxus* (“*S. par*”, strain N17), and the *K. lactis* isolate from an oak in Cheesman Park, Denver (“MMY0329”). Numbers to the left of the gel indicate sizes in Mb of the *S. cerevisiae* chromosomes based on the reference genome (S288C Genome Annotation Release R64.3.1, April 2021). The arrow/cursor icon is irrelevant. (B) Transmitted light micrographs of cells of the Denver oak *K. lactis* isolate MMY0329 following cultivation on rich agar medium (“rich medium”) or 1% potassium acetate (“sporulation medium”). Scale bars are 5 µm. (C) Phylogenetic tree of *K. lactis* isolates from around the world, including the Denver isolate and, as a control, *K. marxianus*, based on *ACT1* sequence. Color-coding is used to label strains according to their published clade assignment based on >100,000 SNPs genome-wide (Friedrich *et al*., 2023). “70-4” is presumed to be the closest relative of the ancestral *K. lactis* strains prior to the acquisition of the *LAC4–LAC12* cluster (Friedrich *et al*., 2023). (D) Sequence alignment of selected nucleotides within *ACT1* comparing the Denver isolate (MMY0329) to sequences from other *K. lactis* populations. Nucleotide “1” corresponds to the 75^th^ nucleotide (inclusive) past the start codon. “•” denotes identity to the first/top sequence. (E) Short-read whole-genome sequence data from the Denver *K. lactis* isolate for the region adjacent to the location of the *LAC4–LAC12* insertion in the dairy reference strain NRRL Y-1140. Colored bars represent individual sequence reads. (E) As in (D) but for a region of the right arm of chromosome 3 showing the presence there of genes C19547, C19519, and C19492. (F) Culture growth, as assessed by absorbance at 595 nm, of cultures of the indicated strains in liquid medium containing either 2% glucose or 10% lactose as the carbon source. MMY0374 is an *S. cerevisiae* isolate from oak 26204 (Table 1). Measurements were made every 5 minutes for 8 replicate cultures. Error bars show the 95% confidence interval. Decrease in absorbance for MMY0329 near the end of the experiment corresponded to clumping of cells around the periphery of the culture well.

To better understand the relationship of our isolate with documented populations, we performed short-read whole-genome sequencing and assembled the genome using either of two reference strains. The assembled genome contained six chromosomes ranging in size from 1.6 to 2.2 Mbp, fully consistent with our PFGE results. *K. lactis* isolates from the wild are typically haploid (Friedrich *et al*., 2023), and the diploid phase is usually short-lived, with meiosis taking place shortly after mating-type switching and mating, all of which are induced by starvation/stress (Herman and Roman, 1966; Barsoum *et al*., 2011). While we did not directly test the ploidy of our isolate, we observed what appeared to be elongated mating cells in addition to asci in a culture transferred to sporulation medium (Figure 3B), which we interpret as evidence that our isolate is haploid and underwent mating type switching and mating prior to sporulation. Due to extra copies of mating type genes present at the silent mating type loci, short-read DNA sequencing cannot determine which allele is present at the mating type locus, and we do not know if the strain is *MAT***a** or *MAT*⍺.

A recent study suggested that polymorphisms in the *ACT1* gene allow powerful discrimination between genetic populations (Lyutova *et al*., 2022), so we constructed a phylogenetic tree based on a larger *ACT1* region from short-read sequences of 41 “domesticated” (dairy) and wild isolates plus the reference strain NRRL Y-1140. This tree almost perfectly recapitulated the relationships described by analysis of >100,000 genome-wide SNPs (Friedrich *et al*., 2023) (Figure 3C), confirming the power of the *ACT1* locus to capture genetic differences. Our isolate did not fit in any of the populations analyzed previously (Figure 4B). We also directly compared our isolate with eight isolates characterized by Sanger sequencing of a portion of the *ACT1* gene (Lyutova *et al*., 2022). Our isolate had a distinct combination of SNPs, notably distinct from the “new” North American population (Figure 3D). No read from our sequencing mapped to the *LAC4–LAC12* gene cluster of the reference strain (Figure 3E). The presence of three genes (C19547, C19519, and C19492) on the right arm of chromosome 3, rather than on the left arm of chromosome 4, further distinguishes our isolate from the *K. lactis* var *drosophilarum* strains that lack *LAC4–LAC12* (Figure 3F). Consistent with the lack of *LAC4–LAC12*, our isolate did not utilize lactose well (Figure 3G). Indeed, our isolate proliferated nearly as quickly as *S. cerevisiae* in glucose-containing medium and nearly as slowly as *S. cerevisiae* in lactose-containing medium, whereas the dairy strain NRRL Y-1140 proliferated at about the same rate in the two sugars (Figure 3G). These data are consistent with a model in which our isolate is the founding member of a previously unstudied genetic population of wild *K. lactis* that is adapted to living on oaks.

**Figure 4.**
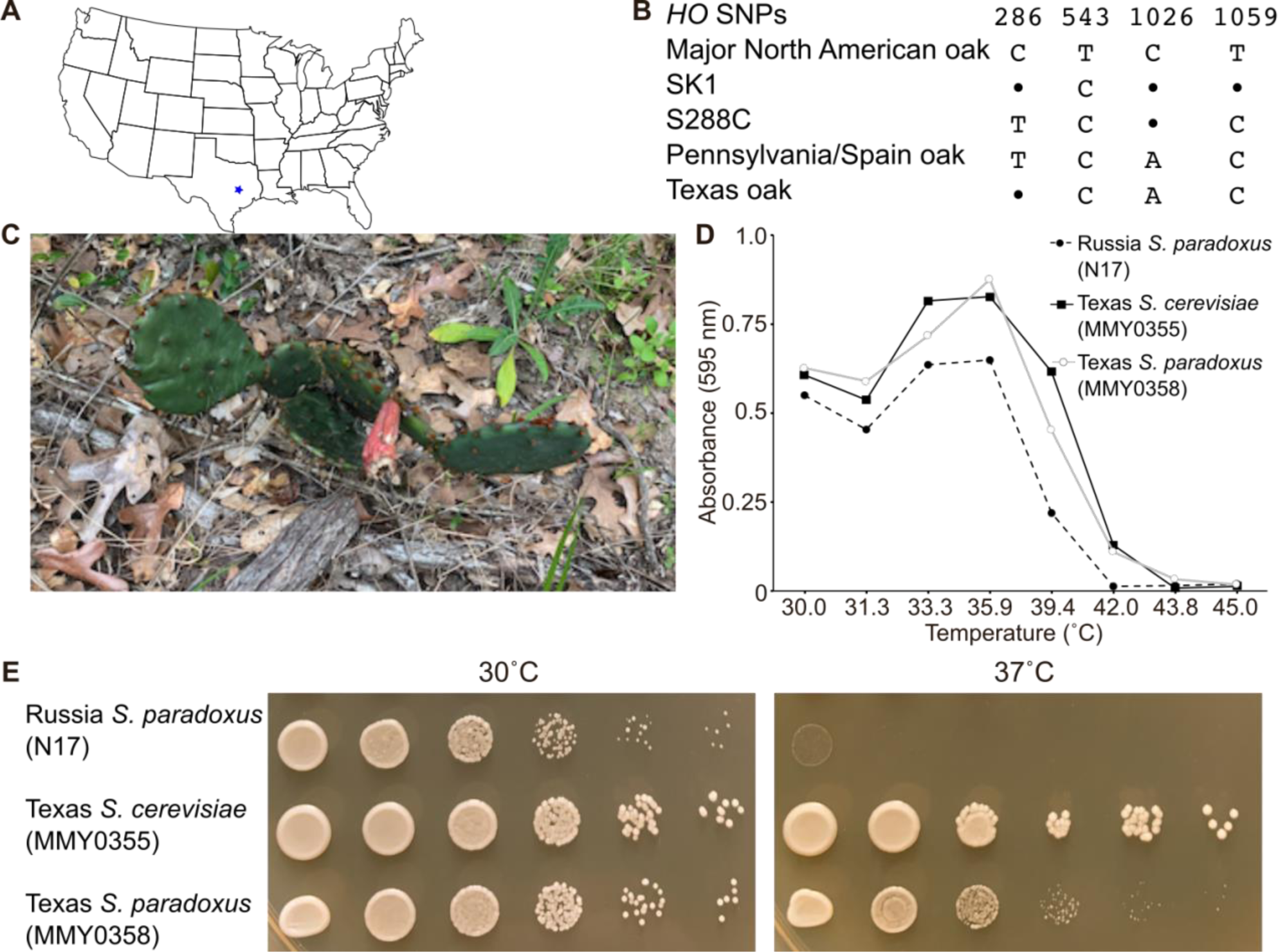
Sampling the diversity within species and genus by varying the geography and ecology. (A) Map of the continental United States showing the location (blue star) in the state of Texas where the oak and sampling was performed. (B) The unique combination of SNPs discovered in the *S. cerevisiae HO* coding sequence from the Texas oak isolates (MMY0351, MMY0355 and MMY0356) is shown by a sequence alignment of selected nucleotides. Nucleotide “1” corresponds to the first nucleotide of the start codon. “•” denotes identity to the first/top sequence, which is the allele found in all our Denver isolates. (C) A photograph of the prickly pear cactus sampled in the Texas oak forest, March 2019. Strain MMY0358 was isolated from the fruit. (D) The growth in liquid rich medium (YPD) at the indicated temperatures was assessed for the indicated strains by measuring the absorbance at 595 nm following ∼21 hr of incubation. (E) The same strains as in (D) were streaked from frozen glycerol stocks to solid rich (YPD) medium, incubated overnight at 30°C, and then cells were scraped from the streaks and resuspended in water. 10-fold serial dilutions were spotted onto new YPD plates with a metal 48-prong replicator and the plates were incubated at the indicated temperature overnight before being photographed.

### A new *HO* allele in *S. cerevisiae* isolated from Texas oaks

By venturing outside of Denver, we found that it was possible to isolate North American oak strains with previously undescribed *HO* sequences. Outside the context of the outreach activity, we used the same isolation method to obtain *S. cerevisiae* from seven post oak trees (*Quercus stellata*) in College Station, Texas (Figure 4A). All the isolates were homothallic. We sequenced *HO* from three of the Texas isolates and found that they were identical, and represented a new allele, encoding the same protein sequence as the other North American oaks, but carrying synonymous SNPs that make it distinct from all others reported here and elsewhere (Figure 4B). The closest available match, differing by only a single SNP, is represented 14 times among the 1,011 isolates, including two isolates from Pennsylvania oaks (CMP, CMQ) and one from an oak in Spain (BBR).

### A heat-tolerant Texan isolate of *Saccharomyces paradoxus*

Interestingly, two of the 14 strains described above (YCX and GAT, which represent the same isolate, originally named UWOPS87-2421) came from a prickly pear in Hawai’i (*Opuntia megacantha*) (Cubillos *et al*., 2009; Liti *et al*., 2009). We noticed prickly pears of undefined species (likely *Opuntia engelmannii*) growing in the same Texas oak forest, and isolated from the fruit of one plant (Figure 4C) a strain that we initially assumed to be *S. cerevisiae*. After failing to amplify *HO* using the same primer pair that was successful with the oak isolates (Figure S2A), we used other primer pairs targeting smaller *HO* fragments and reduced the annealing temperature in the PCR. Two combinations yielded products that were near the expected sizes (Figure S2A), but sequencing revealed that the amplicons instead were 99% identical to portions of the *MHP1* or *FLO11* genes of a strain of *S. paradoxus* (Figure S2B,C). We found presumptive sites of primer annealing to the *S. paradoxus MHP1* and *FLO11* loci (Figure S2D), suggesting that the amplicons arose from spurious primer annealing in the absence of perfect matches to the template genomic DNA. Flo11 (also called Muc1 in *S. cerevisiae*) is a cell surface protein harboring many repeats of a short motif, encoded by tandemly repeated sequences that are prone to recombination-mediated variation in copy number (Verstrepen *et al*., 2005). One presumptive primer binding sequence fell within one such repeat, likely explaining the doublet of PCR products (Figure S2D). To directly test the hypothesis that the prickly pear isolate is *S. paradoxus*, we performed *ITS* sequencing. A 1424-bp *ITS* sequence was a perfect match with *S. paradoxus* (Figure S2E).

We were surprised to find *S. paradoxus* here, where the maximum temperature (41°C during 2012-2019) significantly exceeds 38°C, the suggested maximum for this species (see above). When we compared liquid culture growth in rich medium at different temperatures, the prickly pear isolate of *S. paraxodus* (hereafter MMY0358) was similar to an *S. cerevisiae* strain isolated from a nearby oak tree (Figure 4D). By contrast, an *S. paradoxus* strain (N17) originally isolated from an oak in Russia (Naumov, 1987) had a lower maximum growth temperature (Figure 4D). Plating the same strains on solid rich medium at 37°C confirmed that high temperature tolerance correlated with the climate of origin, not the species (Figure 4E).

## DISCUSSION

Here we describe new biological insights resulting from an activity that engaged children in middle school classrooms in a relatively simple set of hands-on procedures targeted to sixth grade learning objectives in Biology. To broaden the impact of the activity, we also involved university students and PhD students from two university campuses. Moreover, two of the co-authors, N.B. and V.S., designed and executed the grouping analysis of the *HO* sequences from the 1,011 genomes database while they were high school students, as part of the Anschutz Medical Campus CU Science Discovery STEM Research Experience. Thus this activity represents a true partnership spanning multiple ages and levels of education.

The original goal of our activity was to isolate wild *S. cerevisiae* in the hopes of identifying a variety of alleles of *HO*, as was achieved at Mount Carmel (Katz Ezov *et al*., 2010). Taken together with published sequences, however, our results point to limited diversity within North American oak populations. Thus the original puzzle of shared *HO* alleles spread all over the world led to a new puzzle of nearly invariable *HO* sequences across a continent. The Mount Carmel site (dubbed “Evolution Canyon”) features extreme variation in microclimate within a small area, and soil and leaves from many different plants were also sampled to obtain the *S. cerevisiae* isolates (Ezov *et al*., 2006). The limited diversity among North American oak isolates may reflect limited diversity of the ecological niches we sampled in Denver, rather than limited geographical diversity *per se*.

Indeed, by widening the geographical and ecological ranges of our sampling, we broadened the diversity of our isolates. In southeast Texas we discovered a previously unidentified *HO* allele in *S. cerevisiae* living on oaks, and, from a single plant of a different species surrounded by those same oaks, isolated a strain of *S. paradoxus* that has apparently adapted to survive the hot Texas summers. We note that previous studies identified a North American population of *S. paradoxus*, called *SpB*, with distinctly higher maximum growth temperatures than European and other North American populations (Leducq *et al*., 2014), and the *S. paradoxus* strains that best match the *MHP1* and *FLO11* sequences from the Texas isolate belong to the *SpB* population. Furthermore, a very recent study of open agave fermentations also found *S. paradoxus* in parts of Mexico that are similarly hot (Gallegos-Casillas *et al*., 2024). Prickly pears are widespread in the same parts of Mexico, and it is tempting to speculate that the *S. paradoxus* strain we found is part of a heat-tolerant North American population (likely *SpB*) covering a large geographical region. Given this precedent, it should be possible to explore greater *Saccharomyces* diversity in a middle-school outreach context by sampling other plant species, including unroasted coffee and cacao beans of defined origin (Ludlow *et al*., 2016), fruit, and/or flour. The latter could be combined with a sourdough starter activity (Landis *et al*., 2021). Simple phenotypic analysis (e.g. growth temperature) could also be performed in the classroom setting to generate additional useful insights.

We did not originally plan to analyze *K. lactis*, but recent advances in the study of *K. lactis* population genetics (Lyutova *et al*., 2022; Friedrich *et al*., 2023) inspired us to carry out genomic and phenotypic characterization of the single isolate we obtained. Unlike the case with *S. cerevisiae*, our analysis of the *K. lactis* isolate indicates more heterogeneity among North American oak populations for this species. Indeed, based on others’ findings and the hypothesis that in some scenarios *HO* might escape purifying selection (see Introduction), we had hoped to find allele diversity in the *S. cerevisiae HO* gene, but ultimately a *K. lactis* gene (*ACT1*) encoding a highly conserved protein (actin) shows much more variation.

We consider two explanations for this difference that are not mutually exclusive. First, the genes may be fundamentally different, with the allele at *HO* or a nearby gene conferring some functional advantage to *S. cerevisiae* in the oak habitat, whereas *K. lactis ACT1* is a functionally neutral readout of relatedness. However, *S. cerevisiae* oak isolates cluster relatively closely together based on analysis of five (Fay and Benavides, 2005) or 12 (Legras *et al*., 2007) non-*HO* sequences or the entire genome (Liti *et al*., 2009; Peter *et al*., 2018). Furthermore, phylogenetic analysis of *ACT1* sequence alone can recapitulate *S. cerevisiae* relatedness about as well as does *HO* sequence (Figure S3). There is no reason to suspect that *K. lactis ACT1* is under different evolutionary constraints.

Second, the species may be fundamentally different in the mode and extent of their dispersal. *K. lactis* may be more like non-domesticated *Saccharomyces* species (e.g. oak populations of *S. paradoxus*), for which genetic differentiation increases rapidly with physical distance, pointing to limited dispersal (Koufopanou *et al*., 2006; Boynton and Greig, 2014; He *et al*., 2022). By contrast, North American oak populations of *S. cerevisiae* may experience continuous homogenization via rapid/effective dispersal. Indeed, none of the Colorado oaks we sampled is native, and each is <100 years old, hence dispersal must be efficient to populate such new trees. The similarity of North American *S. cerevisiae* oak isolates to each other but not to North American isolates from other natural sources implies that oak strains enjoy a privileged mode of dispersal compared to non-oak *S. cerevisiae* isolates. Considering the very recent isolation of *S. cerevisiae* from butterfly intestines (Bendixsen *et al*., 2021) and of multiple yeast species from the digestive tracts of migrating Taiwanese butterflies (Lin *et al*., 2022), and given the importance of Colorado oaks as butterfly roosts (Scott, 2020), we speculate that migrating insects like butterflies may be important dispersal vectors. It may be possible to include insect sampling in future outreach activities. There may also be strict genetic requirements for stable occupancy of the oak bark habitat, such that non-oak strains arrive just as frequently on oaks but are unable to persist there. Indeed, the fact that we isolated *S. paradoxus* from a cactus within meters of oak trees on which only *S. cerevisiae* was found is probably best explained by the population of *S. paradoxus* in this region of North America being adapted to non-oak niches.

## DATA AVAILABILITY

Strains and plasmids are available upon request. The new *S. cerevisiae HO* sequence (BankIt2837613 Seq PP902360), *S. paradoxus MHP1* (BankIt2842207 Seq PP935650) and *FLO11* sequences (BankIt2842203 Seq PP935649), and the *K. lactis* genome sequence (project PRJNA1128167) have been deposited at NCBI.

Supplemental material is available at XXX.

## ACKNOWLEDGEMENTS

We thank Juan Lucas Argueso for hosting and sponsoring the PFGE analysis; Ken Wolfe for advice about *HO* evolution; Gianni Liti and Gilles Fischer for advice about *HO* sequences in the 1,011 yeast genomes; Bob Sclafani for the NRRLY-1140 strain; Mark Johnston for the N17 strain of *S. paradoxus*; Chris Hittinger for advice on the wild yeast isolation approach and results; Douda Bensasson for the wild yeast isolation protocol and advice; Amy Honors (nee Trujillo) for allowing us to develop the activity in her 6^th^ grade science classroom; Emily Singer, Marc Steingesser, Linnea Wethekam, Lisa Wood, and Katie Alemany for assistance with outreach activities; and the Anschutz Medical Campus STEM Summer Research Experience.

---

**Figure S1.**
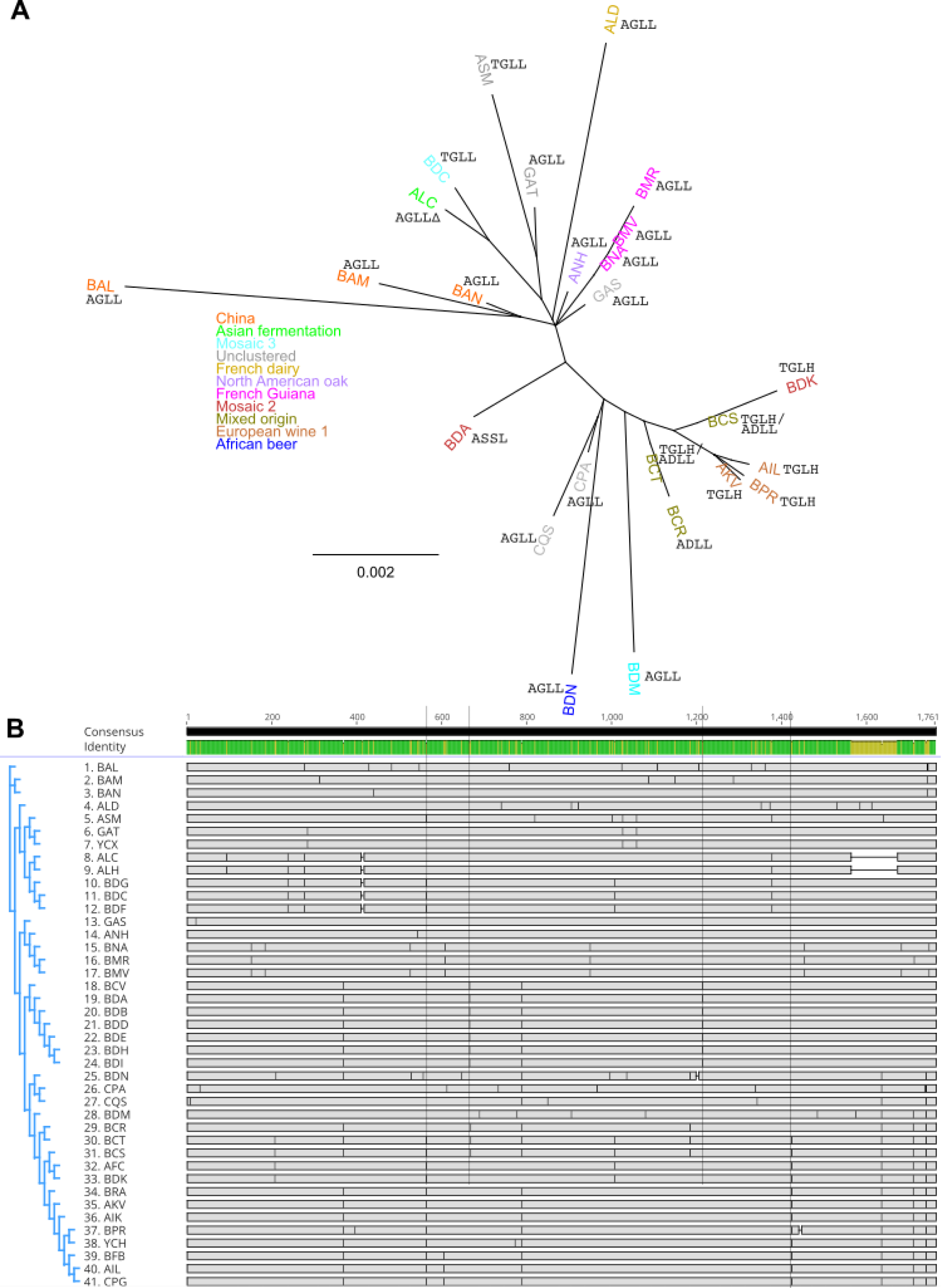
Phylogenetic relationships between natural *S. cerevisiae* isolates based on *HO* gene sequence. (A) Phylogenetic tree of the same 41 isolates mapped in Figure 1B, using the same code to represent the residues at key positions and the presence of the 36-residue deletion. Color coding shows which isolates belong to which clade, as defined in the 1,011 genomes study (Peter *et al*., 2018). (B) Phylogenetic tree and DNA sequence alignment of *HO* coding regions of the same isolates. Vertical lines indicate the positions corresponding to codons 189, 223, 405, and 475.

**Figure S2.**
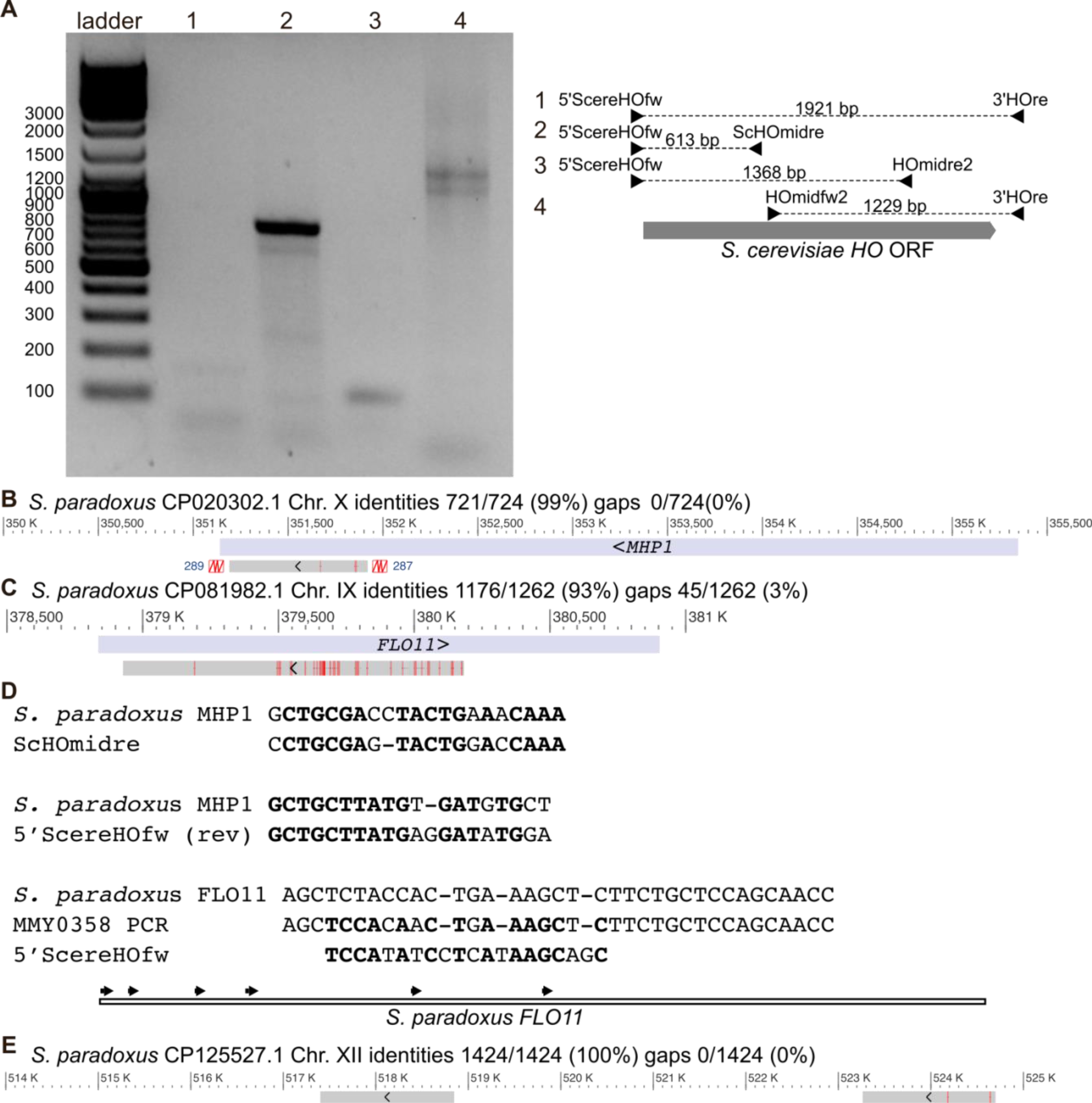
Amplicon sequencing for species identification of the Texas prickly pear isolate. (A) Ethidium bromide-stained DNA following agarose gel electrophoresis of ladder and PCR products generated using the primer combinations shown at right and, as template, genomic DNA from strain MMY0358. The ladder was GeneRuler DNA Ladder Mix (Thermo Scientific #SM0331). The expected amplicon sizes shown at right reflect *S. cerevisiae* sequence. (B) BLAST alignment results for amplicon #2. Red lines indicate mismatches. (C) BLAST alignment results for amplicon #4. (D) Presumptive sites of primer binding in the reference *S. paradoxus* genome. Bold indicates predicted identity with the template. For *FLO11*, the full amplicon was sequenced and is also shown (“MMY0358 PCR”). The extra three nucleotides 5’ of the predicted primer binding site are part of the tandem sequence repeats found in *FLO11* to which the 5’ScereHOfw primer is predicted to anneal. The sequence of one such repeat is shown, and additional copies of this repeat in the sequenced amplicon are shown as arrows in the illustration below. (E) BLAST alignment results for the contig assembled from four *ITS* amplicons (see Methods).

**Figure S3.**
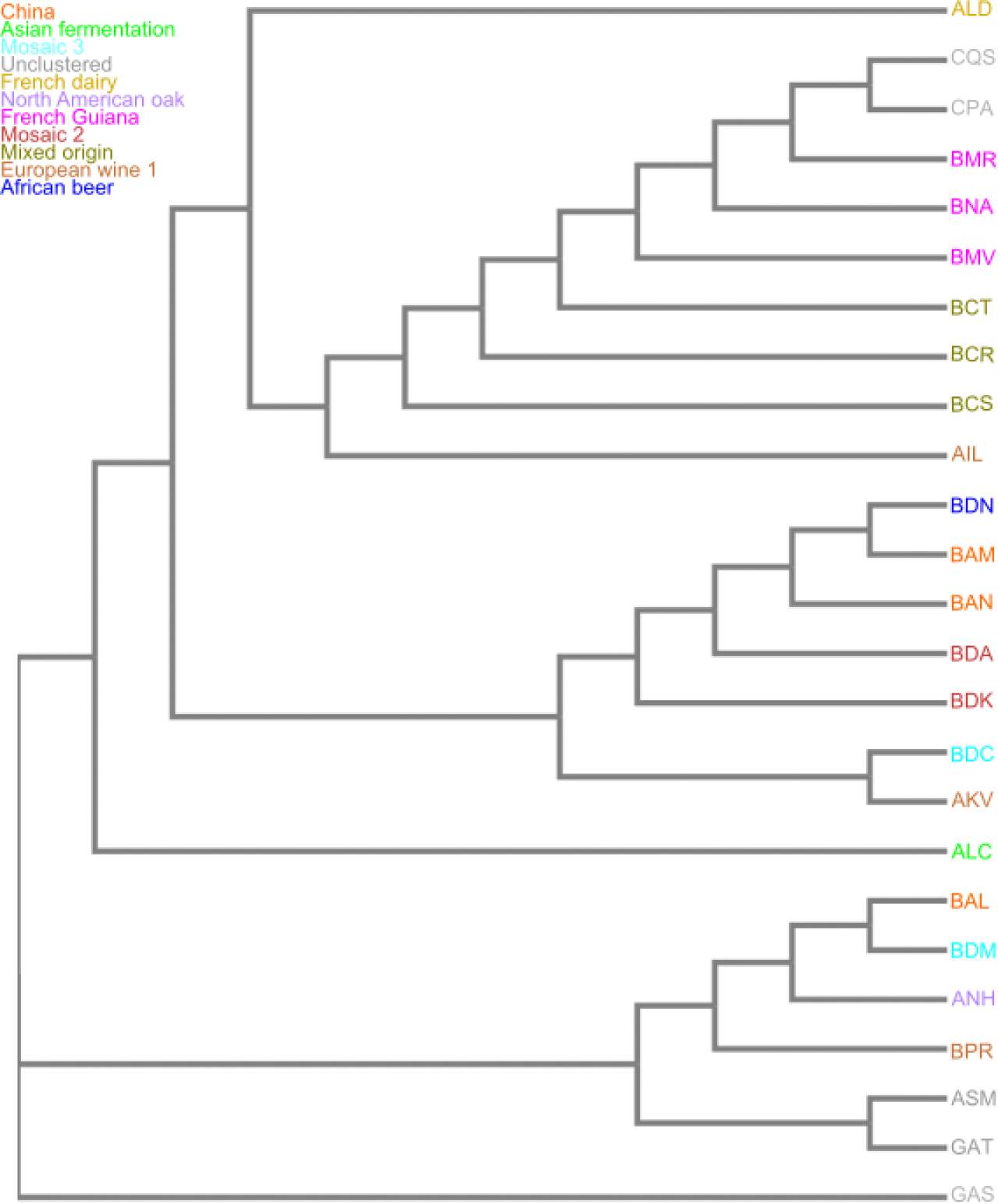
Phylogenetic tree of natural *S. cerevisiae* isolates based on *ACT1* sequence. As in Figure 3C, but with *S. cerevisiae* isolates from Figure 1B and the clade assignments as in Figure S1. Tree was created using the Simple Phylogeny tool at the EMBL-EBI server.

**Table S2.** Connections between lecture topics and Colorado State middle school Science standards and learning expectations.

| Lecture topic | State standard(s)<br>(https://www.cde.state.co.us/apps/standards/7,38,16) | Learning expectation(s) | Science curriculum unit |
| --- | --- | --- | --- |
| *S. cerevisiae* as a microbe and its basic life cycle | All living things are made up of cells, which is the smallest unit that can be said to be alive | Changes in environmental conditions can affect the survival of individual organisms, populations, and entire species | Microbiome, Traits and Reproduction |
|  | Organisms reproduce, either sexually or asexually, and transfer their genetic information to their offspring |  |  |
| Fermentation-focused metabolism of *S. cerevisiae*, natural habitats and industrial use | Organisms and populations of organisms are dependent on their environmental interactions both with other living things and with nonliving factors | Explain and illustrate with examples how living systems interact with the biotic and abiotic environment | Metabolism, Microbiome |
| *HO* gene function in mating-type switching, non-functional *ho* alleles, lack of selection for *HO* when strains don't sporulate | Heredity explains why offspring resemble, but are not identical to, their parents and is a unifying biological principle. Heredity refers to specific mechanisms by which characteristics or traits are passed from one generation to the next via genes | Analyze how various organisms grow, develop, and differentiate during their lifetimes based on an interplay between genetics and their environment | Traits and reproduction |
|  | Genetic variations among individuals in a population give some individuals an advantage in surviving and reproducing in their environment | Explain how biological evolution accounts for the unity and diversity of living organisms |  |
|  | Adaptation by natural selection acting over generations is one important process by which species change over time in response to changes in environmental conditions |  |  |
| Species identification by DNA “barcode” sequencing | Genetic variations among individuals in a population give some individuals an advantage in surviving and reproducing in their environment | Explain how biological evolution accounts for the unity and diversity of living organisms | Traits and reproduction |

